# Motivational Dysregulation with Melanocortin 4 Receptor Haploinsufficiency

**DOI:** 10.1101/2022.05.31.494228

**Authors:** Alex M. Steiner, Robert F. Roscoe, Rosemarie M. Booze, Charles F. Mactutus

**Affiliations:** Cognitive and Neural Science Program, Department of Psychology, University of South Carolina, Columbia, SC 29208

**Keywords:** Melanocortin 4 Receptor, Motivation, Obesity, Stereology, Dopamine

## Abstract

Obesity, by any standard, is a global health crisis. Both genetic and dietary contributions to the development and maintenance of obesity were integral factors of our experimental design. As mutations of the melanocortin 4 receptors (MC4R) are the leading monogenetic cause of obesity, MC4R haploinsufficient rats were fed a range of dietary fat (0-12%) in a longitudinal design. Physiological and motivational assessments were performed using a locomotor task, 5-choice sucrose preference task, an operant task with fixed and progressive ratios, as well as a distraction operant task. Dendritic spine morphology of medium spiny neurons (MSNs) of the nucleus accumbens (NAc), cells with ample D1 and D2 receptors, was also assessed. The percentage of lipid deposits in the liver of each rat was also analyzed using the Area Fraction Fractionator probe for stereological measurements. MC4R haploinsufficiency resulted in a phenotypic resemblance for adult-onset obesity that was exacerbated by the consumption of a high-fat diet. Results from the operant tasks indicate that motivational deficits due to MC4R haploinsufficiency were apparent prior to the onset of obesity and exacerbated by dietary fat consumption after obesity was well established. Moreover, MSN morphology shifted to longer spines with smaller head diameters for the MC4R+/- animals under the high-fat diet, suggesting a potential mechanism for the dysregulation of motivation to work for food. Increasing our knowledge of the neural circuitry/mechanisms responsible for the rewarding properties of food has significant implications for understanding energy balance and the development of obesity.

## 1. Introduction

Obesity, fundamentally a state of excess body fat, occurs because of a prolonged imbalance between the levels of energy intake and expenditure, with the resultant surplus being stored as body lipids (Thomas & Brownell, 2007; Franco, 2009). By any standard, obesity has reached epidemic proportions in the United States; 1/3 of adults are obese and approximately 2/3 of adults are overweight. Over the past 30 years, the prevalence of obesity rose from 7% to 18% in children ages 2 to 19. Further, extreme obesity has reached a prevalence of 5.8% in the same-age children (Ogden *et al*., 2016). The crisis in childhood obesity prevalence foreshadows a further burgeoning in the prevalence of obesity as well as increased risk of morbidity (e.g., diabetes, cardiovascular disease, stroke, etc.)(Thomas & Bronwell, 2007). The prevalence of obesity in the USA has been heralded as having leveled off in the early 2010s (Ogden *et al*., 2016), although the alternative perspective suggests obesity prevalence has consistently risen since 1999, despite a temporary pause in 2009–2012. Indeed, by 2030, 78% of American adults are projected to be overweight or obese (Wang *et al*., 2020). The estimated total national cost of overweight and obese individuals was $149.4 billion yearly, with an average cost of the individual being $1901 yearly (Kim & Basu, 2016). Given that obesity, once established, is so refractory to treatment, longitudinal studies afford an advantageous approach.

Obesity is a multifactorial issue. Both genetic and dietary contributions to the development and maintenance of obesity suggested in human studies have been confirmed in specific animal models (Casper *et al*., 2008; Speakman *et al*., 2008). Analyses of genetic factors of obesity in rodent models have elucidated hypothalamic signaling pathways involved in the control of metabolism and energy balance, the majority investigating deficits in aspects of the leptin signaling pathway. In contrast to the rarity of human leptin deficiency, mutations of the gene encoding MC4R are the most common known form of human monogenic obesity (4–6% of morbidly obese individuals, Vaisse *et al*., 2000; Lubrano-Berthelier *et al*., 2003; a; Farooqi *et al*., 2013). The melanocortin system plays a key role in the central regulation of energy intake, energy expenditure, and energy homeostasis (Ho and Mackenzie 1999; Cone, 2006; Krashes *et al*., 2016). Mutation of the MC4R results in loss of function abolishing correct membrane localization *in vitro*, with *in vivo* data confirming increased body weight, food intake, white adipose mass, and changed substrate preference in male rats (Mul *et al*., 2012). As initially learned from the MC4R mouse knockout, in contrast to most GPCRs, deletion of the MC4R exhibits a gene dosage effect, with heterozygotes exhibiting an intermediate rate of weight gain (Huszar *et al*., 1997). As haploinsufficiency of MC4R is the most common monogenic cause of obesity in humans, the study of melanocortin haploinsufficiency in the trajectory to obesity in the rat was considered of great translational relevance than the study of homozygous knock-outs.

To further our understanding of the behavioral processes and neural mechanisms regulating food reward it is important to consider the neural circuitry that underlies all rewards and motivated behavior, especially given the significant overlap in the underlying pathways (Kenny, 2011; Wise, 2008). To the best of our knowledge, the first experimental test that food may be more reinforcing for obese than lean humans is credited to Epstein and colleagues (Saelens & Epstein, 1996); a replication and extension of which suggested that this effect was particularly noted in obese individuals with the *TaqI* A1 allele (Epstein *et al*., 2007), a putative marker of the DA D_2_ receptor and DA signaling (Jönsson *et al*., 1999; Pohjalainen *et al*., 1998). There is emerging evidence from human pharmacology and/or imaging studies consistent with the postulate of hypofunctioning reward circuitry in obese individuals, with specific alterations in the DA system (Volkow *et al*., 2011). (Evidence for the counter view, that obesity reflects hyperfunctioning reward circuit, cf. Davis *et al*., 2004, does not appear as compelling: although overweight subjects were more hedonic than controls, this pattern reversed among the obese, which were less sensitive to reward; nonetheless, alterations in reward circuitry are implicated). For example, feeding is associated with DA release in the dorsal striatum and the amount of DA release is correlated with meal pleasantness ratings, but not with the desire to eat (hunger) or satiety after eating (Small *et al*., 2003). Further, DA antagonists increase appetite, energy intake, and weight gain, whereas DA agonists reduce energy intake and produce weight loss (de Leon *et al*., 2007; Leddy *et al*., 2004). Dopamine D_2_ receptors are reduced in obese relative to lean individuals and D_2_ receptor measures were negatively correlated with BMI (Wang *et al*., 2001; Volkow *et al*., 2008b). Activation of the striatum in response to highly palatable food is blunted in obese individuals when compared to lean controls (Stice *et al*., 2008). However compelling one may consider the evidence and views about DA hypofunctioning and obesity (e.g., Pijl, 2003; Wang *et al*., 2003; 2004; Volkow *et al*., 2008a; 2008b; Barry *et al*., 2009), it must be recognized that with such cross-sectional clinical data we are at a theoretical impasse. It is unclear whether deficits in reward processing are constitutive and precede obesity, or alternatively, whether excessive consumption of palatable food drives reward dysfunction and thus precipitates diet-induced obesity. The trajectory to obesity begs programmatic study of the MC4R KO rat model of obesity.

The present study investigates the possible physiological and motivational changes due to MC4R haploinsufficiency, dietary fat, or the relationship between the two. Given the known dose-dependent effects of mutation of MC4Rs, the haploinsufficient rat was chosen to model the development of adult-onset obesity. Diets (0% -12% saturated fat) were specifically chosen to be clinically relevant to a range of modern diets. The addition of the inflammatory group allows for a unique control compared to the 12% fat diet group. Consumption of high-fat diets is anticipated to enhance the progression of obesity. Given that 90-95% of cells in the nucleus accumbens are classified as medium spiny neurons (MSNs), cells rich in D1 and D2 receptors that receive dopaminergic signals from the ventral tegmental area (Preston *et al*., 1980), spine morphology in the NAc was also assessed. Our guiding hypothesis is that the trajectory to obesity is preceded by alterations in motivational systems, including neuroadaptations in the central nervous system; these alterations in motivational systems will have persistent functional consequences for vulnerability to excessive caloric intake in an obesogenic environment, and the extent of central nervous system neuroadaptations will be exacerbated in an obesogenic environment.

## 2. Methods

### 2.1 Subjects

Both normal and mutant (MC4R +/-) male Wistar rats (ns=33) were procured (Transposagen, Lexington, KY) and weaned at postnatal day 21. Originally, female wild-type Wistar P generation rats were bred with MC4R -/- male rats, resulting in the MC4R +/- F1 generation; this specific breeding approach was performed to control for any potential fetal development confound of altered maternal care by a genetically modified female. After weaning, animals were housed in pairs with one haploinsufficient rat and one control rat per cage (which were fed the same diet, as indicated below in experimental design). Animals were kept in an AAALAC accredited facility at 21 ±2ºC, 50% ± 10% relative humidity on a 12-hour light/dark cycle with lights on at 07:00h EST. All behavioral testing was conducted during the light cycle. The Institutional Animal Care and Use Committee (IACUC) of the University of South Carolina approved the project protocol under federal assurance (#D16-00028).

### 2.2 Experimental Design

The MC4R +/- rats and wild-type control pairs in each residential cage were randomly assigned to one of four diet groups (Modified AIN-76 diets, Bio-Serve, Frenchtown, NJ), thus constituting a 2 × 4 factorial design. Specifically, the diets included a control diet (n=9 per group) (1.7% Saturated Fatty Acids SFA, with 12.2% total kcal from fatty acids), a low-saturated-fat diet (n=8 per group) (6% SFA, with 40% total kcal from fatty acids), and a high-saturated fat diet (n=8 per group) (12% SFA, with 40% total kcal from fatty acids). The fourth group was an inflammatory diet matched to the 12% high-fat diet group (n=8 per group) (1.7% SFA, with 12.2% total kcal from fatty acids, 20:1 ratio of omega-6:omega-3 unsaturated fatty acids). While in their home cages, animals had *ab libitum* access to food and water. Diets were chosen to replicate a range of possible diets relevant to human consumption.

### 2.3 Body Measurements

Body weights, crown-rump length, and waist circumference were taken on postnatal days 21-23, 27-29, 34-36, 41-43, 48-50, 62-64, 76-78, 90-92, 120-122, 152-154, and day of sacrifice. BMI was employed as an estimate of obesity (Novelli *et al*., 2007) and calculated as body weight (g)/(body length (cm)^2^. One MC4R +/- KO animal on the INF diet was found deceased on PD 98; missing data for this animal was replaced via imputation. One MC4R +/- on the CON diet died prior to the variable PR test, and n(s) were modified accordingly.

### 2.4 Timeline of Behavioral and Motivational Assessments

Activity tasks began for all animals at day 30 days of age. All animals repeated locomotor activity and sucrose preference tasks throughout a significant portion of their lifespan. Prior to the development of obesity, animals were assessed using fixed ratio and progressive ratio operant tasks to assess motivation. Following the onset of obesity, motivation was assessed similarly, using variable progressive ratio and distraction operant tasks. Post sacrifice, dendritic spine analysis, and liver steatosis analysis occurred. The overall study design can be seen in Table 1.

**Table 1.** Timeline for Dependent Measures

| Activity Tasks. Postnatal days 30, 60, 90, 120, 150, and 180 |  |  |
| --- | --- | --- |
| Sucrose Preference Task |  | Locomotor Activity |
| Pre-Obesity Motivational Tasks. Postnatal days 61-120 |  |  |
| Fixed Ratio 1, 3, 5 |  | Progressive Ratio |
| Post-Obesity Motivational Tasks. Postnatal day 120-260 |  |  |
| Variable Progressive Ratio | No Distraction Task | Distraction Task |
| Sacrifice |  |  |
| Dendritic Spine Analysis |  | Steatosis Analysis |

### 2.5 Locomotor Activity

The testing apparatus for the locomotor activity task was a 40 cm by 40 cm square chamber with a circular Plexiglas insert to promote movement. The chamber tracks ambulation and rearing using infrared photocells on X, Y, and Z dimensions (Hamilton-Kinder Inc., Ponway, CA). Photocells were tuned by the manufacturer to control for the extra Plexiglas insert. The test was administered at postnatal days of age 21, 30, 60, 90, 120, 150, and 180 under low light conditions to simulate the nocturnal experience when rats are active. The beam breaks across the photocell grid (32 × 32, spaced 2.5 cm apart) were recorded using Digipro System Software (v. 140, AccuScan Instruments) in real-time. Motor Monitor software (Hamilton-Kinder Inc, Ponway, CA) was used to record and monitor movements inside the chamber. Basic movements were defined as a clearing of a beam when a new beam is broken, while rearing was defined as a breaking of an overhead beam.

### 2.6 Sucrose Preference Test

Sucrose preference testing was administered on days 30, 60, 90, 120, 150, and 180. The animals were habituated to the testing cage on postnatal day 21. For a 20-minute testing session, five sucrose solutions (0, 1, 3, 10, and 30% by volume) were available to the animal. Bottle weight differences were used for the preference analysis. Potential position preference was controlled by using block-randomization and the Latin-square procedure for bottle sequence.

### 2.7 Operant Testing Apparatus

The operant task chambers (ENV-008; MED Associates, St. Albans, VT) were housed in a sound-attenuated cabinet. The front of the chamber had access to a recessed dipper through a 5 cm by 5 cm window with infrared sensors to track nose poke time in seconds. The dipper has a 0.1 ml cup attached, which was raised into the chamber to allow access to the cup. The cup contained a sucrose solution upon the completion of the required responses. On each side of the opening, two retractable metal levers were located 7.3 cm above the metal grid floor. On the back wall of the apparatus was a third, inactive, lever also located 7.3cm above the metal grid floor. At the beginning of testing, all three levers were available. Animals underwent various ratio schedules to learn to respond for continuous reinforcement during 82-minute sessions. After correct operant responses to the active lever, the sucrose reward solution was presented for 4 seconds, whereas responding on the inactive lever was recorded but not reinforced.

### 2.8 Fixed-Ratio and Progressive-Ratio Tasks

On Postnatal day 61, animals underwent a fixed-ratio (FR) 1 schedule for at least 3 days. After three consecutive days of stable responding, defined by greater than 60 rewards during the test period, the animals were moved to an FR-3 schedule. Similarly, after 3 consecutive days of stable responding, defined by 120 rewards on the FR-3 schedule, animals were moved to an FR-5 schedule. Upon 3 consecutive days of stable responding on the FR-5 schedule, animals underwent a progressive ratio test. The sequence of lever pressing requirements were 1, 2, 4, 6, 9, 12, 15, 20, 25, 32, 40, 50, 62, 77, 95, 118, 145, 179, 603, 737, 901, 1102, 1347, 1646, and 2012, for a maximum of two hours in length for each test. The use of a geometric ratio (defined as *nj*=5*e*j/5-5) is well suited for the examination of satiety, as the requirements for response increase exponentially after each reinforcement (Richardson & Roberts, 1996; Killeen *et al*., 2009).

### 2.9 Variable Progressive-Ratio task

On postnatal day 220, animals underwent the same progressive ratio schedule, with varying concentrations of sucrose reward (1%, 3%, 5%, 10%, or 30%). Each animal received a test for each concentration with a 5% sucrose FR-5 schedule on days in between tests. The total testing took 10 days with a 5% sucrose concentration PR schedule on the last day to prevent extinction. Starting concentrations were block randomized with concentrations shifted according to a Latin-square design

### 2.10 Distraction Task

Upon completion of the progressive ratio task at approximately 230 days of age, animals performed an FR-5 schedule distraction task for 60 minutes. The first test was an FR-5 schedule with a distracting tone (5dB above the background fan noise of the chamber) during the middle 20 minutes of the 60-minute test period. The next day, animals were placed on an FR-5 schedule again, with no distraction. Lastly, on the third day of testing, animals were tested on the same FR-5 schedule with the distracting tone played from minutes 5-25, with no tone being played during the remaining of the testing period.

### 2.11 Preparation of Dendritic Spine Analysis

Medium spiny neurons, cells with ample D1 and D2 receptors, represent the majority of the cellular makeup of the nucleus accumbens and network to the ventral tegmental area (Preston *et al*., 1980). Synaptodendritic alterations on the medium spiny neurons as measured by changes in spine morphology may contribute to or underlie neurobehavioral responses to genotypic or dietary modifications. Following completion of the distraction task, animals were sacrificed throughout PD266-PD282.

Animals were sacrificed by first being anesthetized using sevoflurane (Abbot Laboratories, North Chicago IL), followed by transcardial perfusion with 100 ml of 100 mM PBS wash then by 200-250 ml of 4% paraformaldehyde buffered in PBS (Sigma-Aldrich, St. Louis, MO). Brains were then extracted and fixed in 4% paraformaldehyde until further processing. Using a rat brain matrix (ASI Instruments, Warren, MI), 200µm thick coronal slices were cut, washed in PBS 3 times, and placed in tissue cell culture plates (24 well; Corning, Tewksbury MA) for DiOlistic Labeling.

DiOlistic Cartridges were prepared by first dissolving 300mg of tungsten beads (Bio-Rad, Hercules, CA) in 99.5% pure methylene Chloride (Sigma-Aldrich, St. Louis, MO) and sonicated in a water bath (Fisher Scientific Fs3, Pittsburgh, PA) for 30-60 minutes. Crystallized DiI (14.5mg; Invitrogen, Carlsbad, CA) was dissolved in methylene chloride and then protected from light until further application. Upon completion of the sonication, 100 µl of the bead solution was placed on a glass slide and titrated with 150 µl of the DiI solution. The solutions were slowly mixed using a pipette tip and then allowed to air dry. The mixture was then transferred to a 15 ml conical tube (BD Falcon, San Jose, CA) along with 3 ml ddH2O and sonicated in a water bath for another 30-60 minutes.

Tefzel tubing (IDEX Health Sciences, Oak Harbor, WA) was cut into 1.7 M lengths to match the length of the tubing preparation station (Bio-Rad, Hercules, CA). Using 10ml ddH2O, 100 mg of polyvinylpyrrolidone (PVP; Sigma-Aldrich, St. Louis, MO) was dissolved and vortexed. The solution was passed through the length of the tubing to aid in bullet adhesion. The 3 ml bead and dye solution was drawn into the tubing and allowed to spin for 5 minutes to uniformly coat the tube. The tube was then allowed to dry spin for approximately 10 minutes with a nitrogen gas flow of 0.1 LPM, followed by an adjustment to 0.5 LPM, and spun for another 60 minutes to ensure a dry tube. Tubing was then cut into 13 mm segments and stored in dark until use.

### 2.12 DiOlistic Labeling and Medium Spiny Neuron Quantification

Bullets were loaded into a Helios gene gun (Bio-Rad, Hercules, CA), with He gas flow adjusted to 90 PSI and 3 µm pore filter paper (Millipore, Billerica, MA). The gene gun barrel was placed approximately 2.5 cm away from the sample and dye was delivered directly onto the slice. Tissues were then washed with PBS and stored overnight at 4°C. The following day tissues were mounted using Pro-Long Gold Antifade (Invitrogen, Carlsbad, CA), coverslipped (ThermoFisher Scientific, Waltham, MA), and stored in the dark until examined via confocal microscopy.

Medium spiny neurons (MSNs) from the nucleus accumbens (NAc), identified by Bregma’s landmark using a rat brain matrix (Paxinos & Watson, 2014), were used for the analysis. Using a Nikon TE-2000E confocal microscope with Nikon’s EZ-CC1 software (version 3.81b), Z-stack images were obtained. DiI fluorophore was excited using a green helium-neon (HeNe) laser with an emission of 533 nm. Images were captured using a 60X objective with a numerical aperture of 1.4. Z plane intervals were 0.15-0.30 µm (pinhole size 30 µm) with an internal 1.5X additional magnification. Once images were captured, morphometric analysis of spines was performed using Neurolucida (version 11.01.1), utilizing the AutoNeuron and AutoSpine extension modules (MicroBrightfield, Williston, VT).

Length, volume, and head diameter parameters of dendritic spines were used for analysis. Spine lengths were defined as between 0.01 µm to 45 µm (Blanpied & Ehlers, 2004). Spine head diameters were defined as between 0.3 µm and 1.2 µm (Bae *et al*., 2012). Lastly, volume parameters were defined as between 0.02 µm^3^ and 0.2 µm^3^ (Merino-Serrais *et al*., 2013).

### 2.13 Steatosis Analysis

Livers of all animals were extracted and stored in a -80°C freezer until processed. Livers (N=32; n=4 per group) were randomly selected to undergo stereology procedures. Each liver was sectioned into 20-micron slices using a cryostat (Shandon Cryotome). Every 18^th^ slice was mounted and subsequently stained for lipid deposits using Oil Red O. The following was the histological staining procedure:

1. Slices were mounted and placed in a 10% PFA solution for 8-10 minutes
2. Washed with distilled water
3. Placed in 100% propylene glycol for 3-5 minutes
4. Placed in Oil Red O heated to 60ºC for 8-10 minutes
5. Placed in an 85% propylene glycol and distilled water solution
6. Finally washed once more with distilled water.

To estimate the percent volume of fat in each liver a Nikon Eclipse E800 (Nikon, Melville, NY) equipped with a motorized LEP MAC 5000 XYZ stage (Ludl Electronic Products, NY) and Stereoinvestigator (MicroBrightfield Williston, VT, Version 11.09) were used. The Area Fraction Fractionator probe allows randomly selected sampling sites to be determined and used to estimate volume with a sampling grid (West, 2012). For each slice, four sampling sites were determined with a 200×200 µm square with markers 8 µm apart (a total of 625 markers) laid over each sampling site. From the stereological count, an accurate estimation of the percent volume of fat was calculated by taking points counted divided by total points.

### 2.14 Statistical analysis

All Statistical analyses were done using IBM SPSS v 24 (IBM Corp., Somers, NY). Graphs and curve fits were made using GraphPad Prism 5.02 (GraphPad Software, Inc. La Jolla, CA). On postnatal day 98, one MC4R +/- haploinsufficient animal on the inflammatory diet was found deceased. Missing data for the animal was replaced by imputation. To detect if there was an effect of litter, we conducted a repeated-measures ANOVA on the bodyweight data using litter and genetic condition as variables (McLaurin & Mactutus, 2015; McLaurin *et al*., 2020). As the factor of litter was found non-significant at this alpha level [F(8,56)=1.78, P=0.10], statistical analysis proceeded independent of litter.

BMI was analyzed using a mixed-model ANOVA with genetic condition and diet as between-subjects factors and time (day) as a within-subject factor. Two separate mixed-model ANOVAs were run for basic movement and rearing during the locomotor activity tasks. Similarly, condition and diet were between-subject factors whereas time was a within-subject factor. The sucrose preference task data was also analyzed using a mixed-model ANOVA. The same factors as the previous analyses were used, as well as the addition of the within-subject factor of sucrose concentration.

The progressive ratio task was analyzed using a simple between-subjects ANOVA using genetic condition and diet. The variable progressive-ratio task was analyzed using mixed-model ANOVAs using the same factors as the progressive-ratio in addition to sucrose concentration as a within-subjects factor. The distraction task was analyzed using mixed-model ANOVAs as well. The within-subjects factor for the analysis was the 5-minute bins that were recorded throughout the task.

Liver steatosis was analyzed using a between-subjects ANOVA.

Potential shifts in the parameters of spine morphology were assessed with an analysis of the entire distribution of both spine length and head diameter.

## 3. Results

### 3.1 Both MC4R +/- haploinsufficiency and consumption of high-fat diet cause obesity

BMI data were used to assess the effect of MC4R +/- (Figure 1). Both genetic condition [F(1,26)=25.50, P≤.001] as well as diet [F(3,26)=3.84, P≤.05] exerted statistically significant effects on BMI. The three-way interaction of genetic and diet conditions with age indicated differential growth as a function of genetic condition that was modified by diet. Figure 1 depicts the differences between both the MC4R +/- and background strain [F(1,401) = 14.93, P≤.001] as well as the difference between the 0% and 12% high saturated fat diet groups [F(1,203) = 37.82, P≤.001]. Most interestingly, there are notable differences in weight as a function of MC4R haploinsufficiency at approximately 60 days of age and as a function of the obesogenic diet at approximately 120 days of age, consistent with the development of adult-onset obesity.

**Figure 1:**
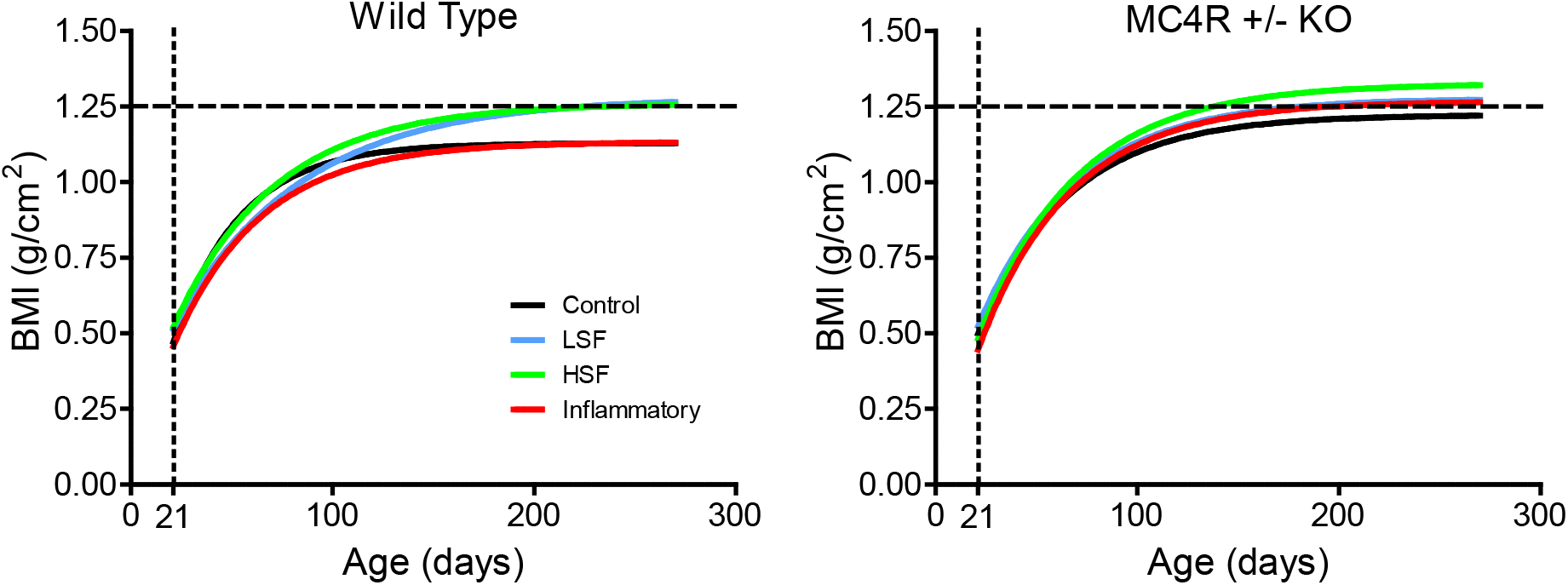
Bodyweight of both control and MC4R +/- animals. (A) Haploinsufficient animals show a significantly higher plateau than their control counterparts. Divergence of growth curves is apparent in young adulthood, at approximately 60 days of age. (B) Under an obesogenic environment, there was a significant difference between the high-fat and control diets. Divergence of growth curves is apparent at approximately 120 days of age.

### 3.2 MC4R +/- haploinsufficiency and obesogenic environment influence rearing behavior, but not overall locomotor activity

Locomotor activity was analyzed by measuring the area under the curve across age for each genetic group and dietary condition for both dependent measures, as depicted in figure 2. Basic movement measures showed no difference between genetic groups (global curve fit) but suggested a slight decline with an increase in dietary fat. For rearing, control and MC4R +/- animals yield divergent outcomes. As dietary fat increases, control animals yield an increase in rearing. Conversely, the MC4R +/- animals yield a decrease in rearing as dietary fat increases. Control and MC4R +/- animal curve fits for the rearing were 0.92 and 0.98, respectively, with a pronounced difference between the two lines [F(2,2)=62.17, P≤0.016].

**Figure 2:**
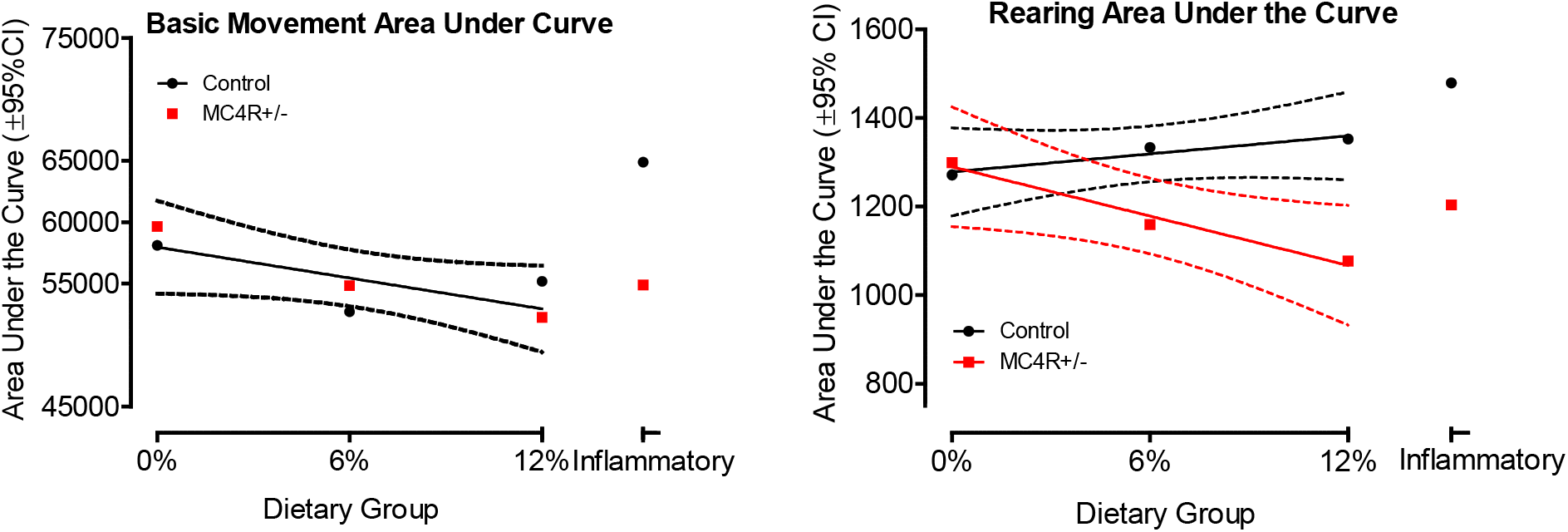
Area under the curve measures for both basic movement and rearing were fit to curves of activity across the test ages. Basic movement yielded no statistically significant difference between genetic groups as a function of dietary fat. However, rearing behavior displayed divergent results for control and MC4R +/- animals as a function of dietary fat [F(2,2) = 62.17, P≤0.016].

### 3.3 Sucrose preference is altered by MC4R +/- mutation as well as dietary fat

The 5-choice sucrose preference test revealed an altered searching and sampling pattern as a function of either dietary fat or MC4R haploinsufficiency, as displayed in figure 3. As early as 60 days of age, there was a striking increased preference for lower sucrose concentrations among Wistar rats fed the control versus high-fat diet as well as by MC4R +/- animals relative to Wistar controls. At 6 months of age, the concentration preference curves for the MC4R +/- groups displayed the most consistent dose-response effects relative to those of the Wistar background control animals, suggesting an increased sensitivity to manipulation of sucrose concentration.

**Figure 3:**
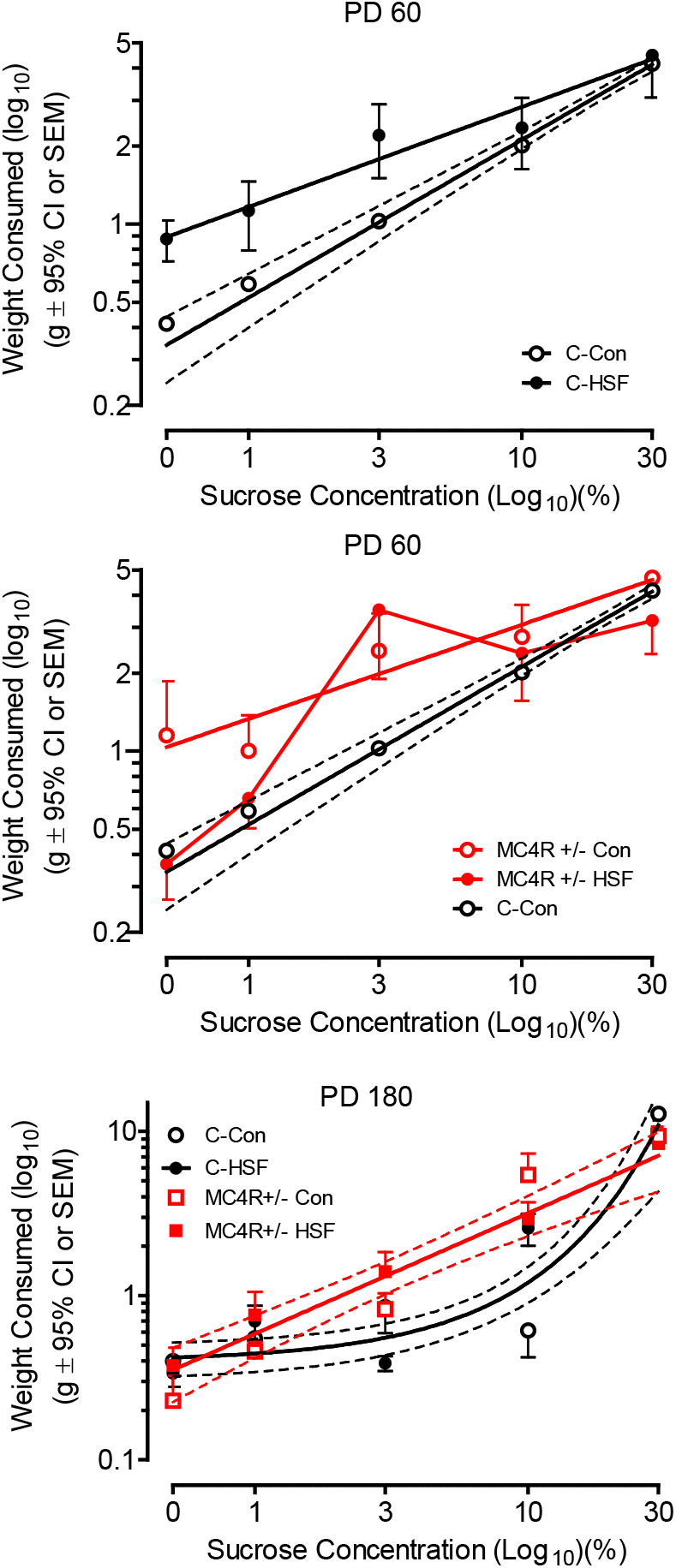
MC4R +/- KO rats exhibit an altered searching pattern in a 5-choice sucrose preference test. Tests performed on PD 60 (A-B) and PD 180 (C) are illustrated. (A) Alterations of preference are apparent as a function of sucrose concentration comparing control and high saturated fat diet groups [different slopes, F(1,6) = 19.69, P≤0.005]. (B) MC4R+/- animals displayed a greater preference for low sucrose concentrations than control animals [different slopes, F(1,6) = 9.15 P≤0.025]. (C) The slope of the concentration preference curve for the MC4R +/- groups displayed a prominent linear dose-response effect (global curve fit, r^2^=0.91) whereas the concentration curves for the genetic background control animals displayed an exponential growth function (global curve fit r^2^=0.65) with sensitivity to only the highest sucrose concentrations.

### 3.4 Motivational deficits are present early in the trajectory to obesity

The fixed-ratio and progressive-ratio operant tasks were used to analyze motivational differences. The tasks were conducted prior to the onset of obesity. None of the fixed ratio operant tasks (schedules FR1, FR3, and FR5) revealed a significant effect of either genetic condition or diet (data not shown).

The progressive-ratio operant task that was assessed, beginning at postnatal day 105, illustrated that the MC4R+/- animals displayed significantly superior performance relative to Wistar control animals on each of the four measures recorded. The increased responding of the MC4R +/- group [F(1,16)= 11.65, P≤0.05] is shown in figure 4.

**Figure 4:**
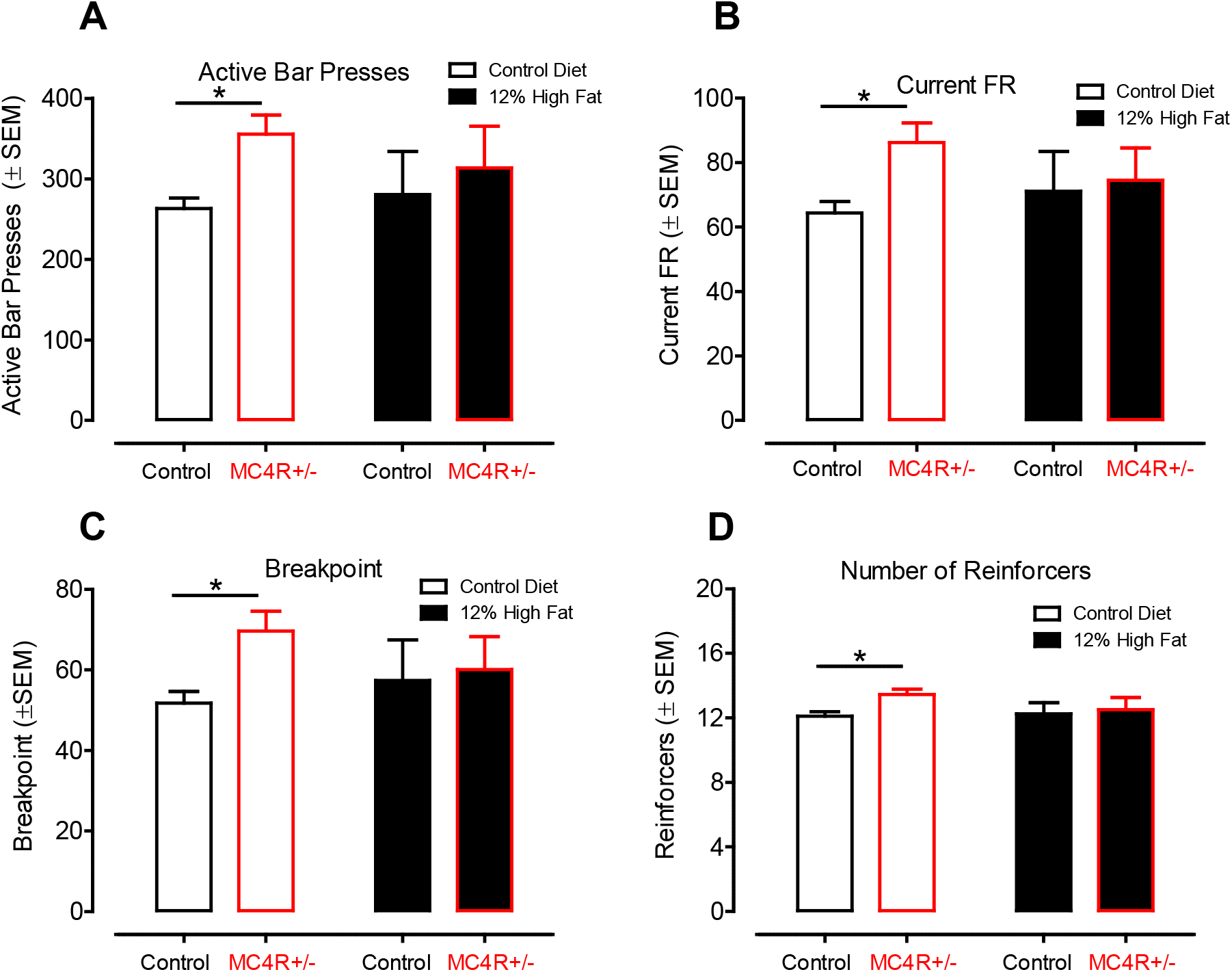
Results from the progressive ratio task started at postnatal day 105. MC4R+/- animals fed the control diet show a clear increase in performance across all four measures indicating increased motivation for food compared to their control counterparts. The same task with animals fed the high saturated fat diet failed to detect any difference between the control and MC4R+/- groups. A: Active lever presses. B: Current fixed-ratio schedule. C: Breakpoint. D: Number of reinforcers received.

### 3.4 After obesity is well established, motivational regulation is dependent on dietary fat

The variable progressive-ratio operant task was assessed at postnatal day 220 to investigate motivational differences with varied reward concentrations well after obesity was established. The factor of dietary fat was found statistically significant [F(3,283)=4.27, P≤.05]. More importantly, the interaction between MC4R mutation and diet was statistically significant [F(3,283)=2.63, P≤.05]. Animals fed the control diet showed a similar increase in responding with an increase in sucrose concentration, regardless of their genetic condition. MC4R+/- animals fed the high saturated fat diet show increased responding regardless of the sucrose concentration reward. The control counterparts only reach similar responding levels with the highly rewarding 30% sucrose concentration. The effect of diet and genetic condition on the variable-ratio task can be seen in figure 5.

**Figure 5:**
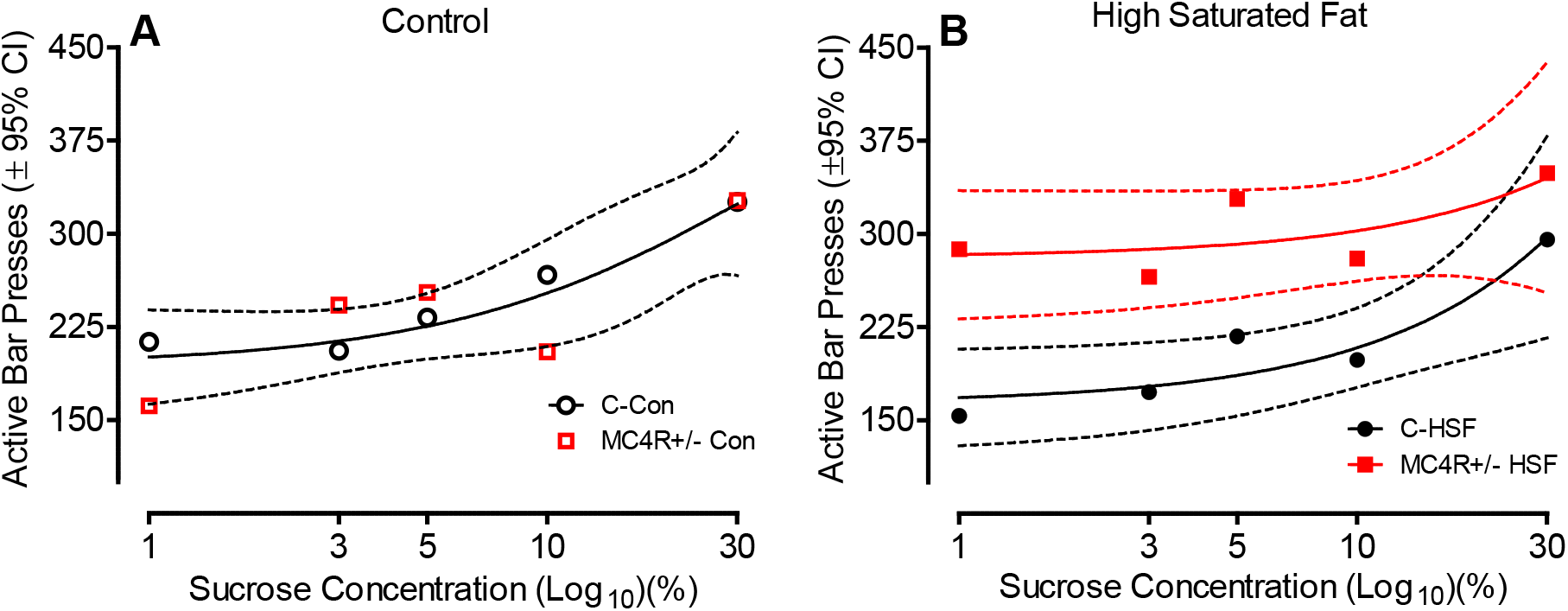
Results from the variable progressive ratio task starting at postnatal day 220, separated by dietary group. (A) Animals fed the control diet do not show any difference in bar presses as a function of genetic group; both groups increased performance as a function of sucrose concentration (one-phase association, global curve fit r^2^ = 0.75). (B) MC4R+/- animals fed the high saturated fat diet show an increase in responding for a lower sucrose concentration than the control counterparts, however, at a high sucrose concentration responding rates for both groups indicate no difference [(F2,6) = 19.67, P≤0.002].

The no distraction and distraction task data collected at 7-8 months of age is illustrated in figure 6. The FR5 schedule used in the no distraction task, as analyzed by ANOVA, revealed a significant effect of diet condition [F(7,88)=5.70, P≤0.05]. Performing a Tukey’s post hoc analysis revealed a striking and significant increase in rewards earned by the MC4R rats relative to Wistar controls under the obesogenic environment provided by the high saturated fat diet; no differences in performance were observed under the control diet condition.

**Figure 6:**
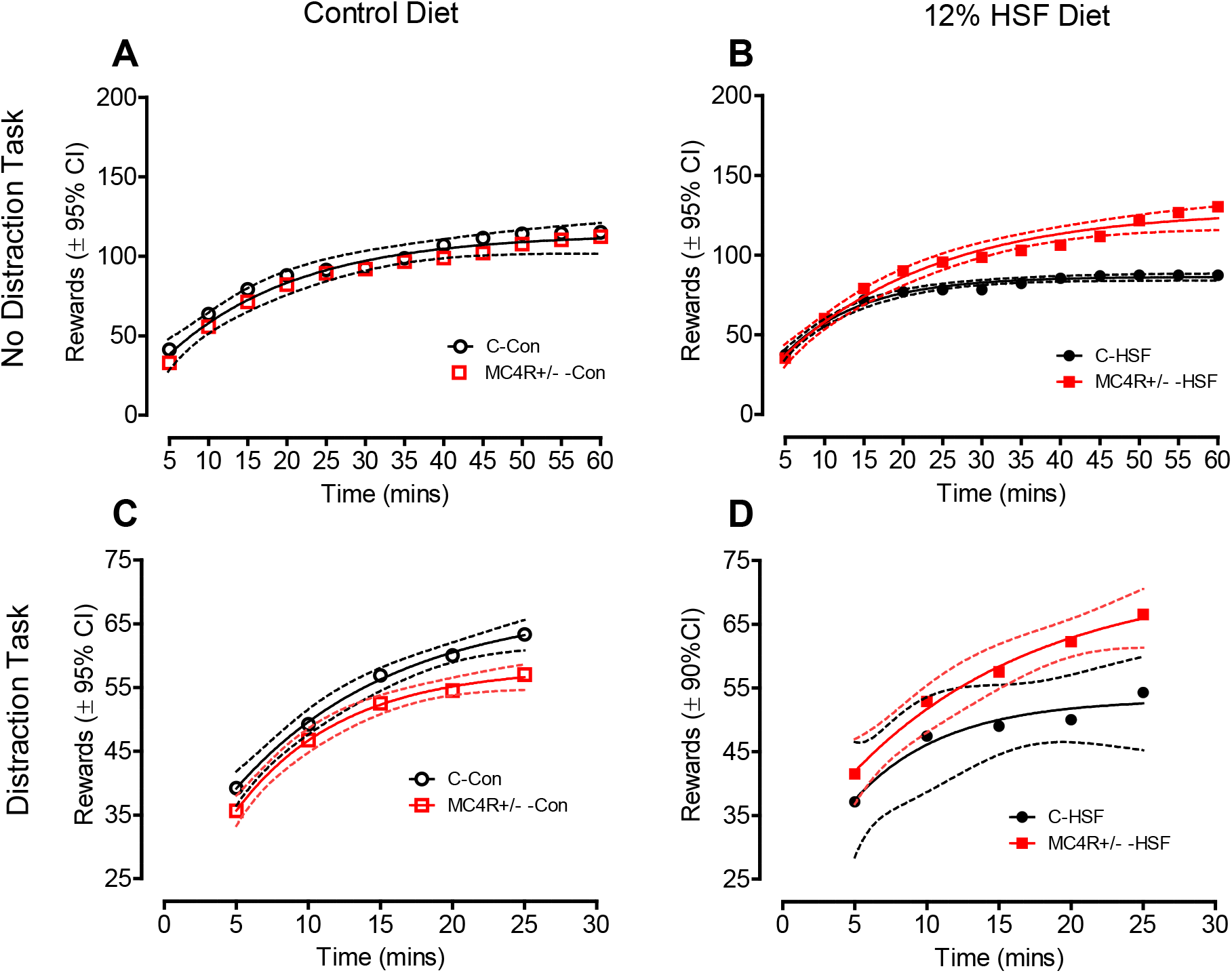
Distraction and no distraction FR5 tasks starting on postnatal day 230. (A) During the no distraction task, animals fed the control diet did not differ in rates of responding as a function of the MC4R mutation. (B) MC4R +/- KO animals fed the high saturated fat diet respond at significantly higher rates than their control counterparts. (C) The presence of a distracting tone disrupted rewards earned by the MC4R +/- animals relative to those earned by the controls. (D) MC4R +/- rats fed the high saturated fat diet were resistant to the distracting tone earning significantly more rewards than the controls animals fed the high saturated fat diet.

With the presentation of a distraction tone, significant alterations in the performance of the MC4R +/- animals were observed as a function of diet condition. Under the control diet, the MC4R +/- rats earned fewer rewards than Wistar controls whereas under the high saturated fat diet the MC4R +/- rats displayed significant resistance to the disrupting effect of the tone earning significantly more rewards than the Wistar controls.

### 3.6 Medium Spiny Neurons morphology shifts in MC4R+/- animals fed a high saturated fat diet

Analyzing MSN spine data we see a population shift in both the diameter and length in MC4R+/- animals fed the high saturated fat diet, as illustrated in figure 7. Compared to controls that displayed relatively short-length spines, haploinsufficient animals demonstrated a relative population shift to longer spines [(χ^2^(18)=45.92, P≤0.001]. Regarding head diameter, the controls displayed relatively greater diameter spine heads whereas the haploinsufficient animals demonstrated a population shift to smaller spine head diameters [(χ^2^ (16)=72.96, P≤0.001].

**Figure 7:**
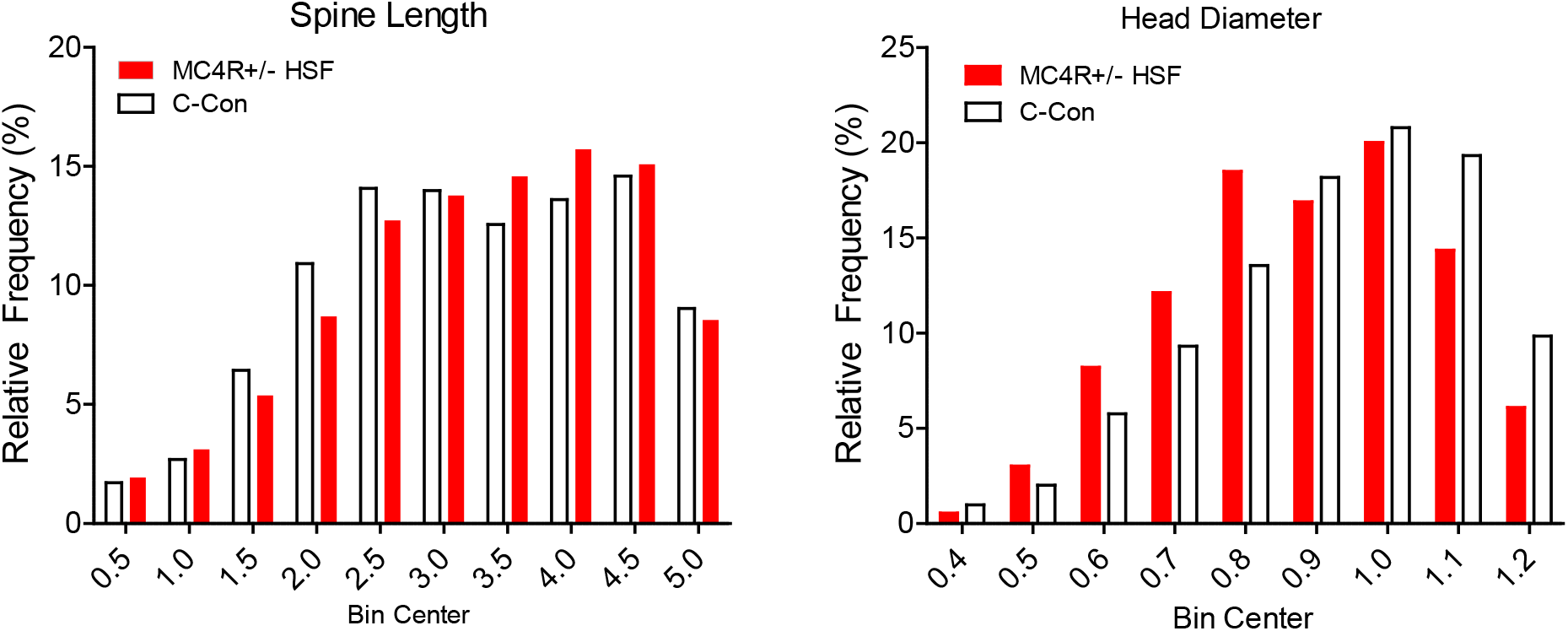
Analysis of MSN spine length (A) and head diameter (B) both displayed a population shift due to the effect of MC4R haploinsufficiency as well as a high saturated fat diet. MC4R+/- animals fed the high saturated fat diet have a higher population in morphologically longer spines with smaller head diameters relative to controls with shorter spines with larger head diameters.

### 3.5 Development of steatosis is linearly dependent on dietary fat consumption

The steatosis analysis found a significant effect of diet [F(1,3)=5.40, p≤.05], as illustrated in figure 8. As may be observed, the accumulation of fat in the liver had a direct linear relationship to the percentage of fat in the diet. Specifically, dietary fat was predictive of the degree of liver steatosis (r^2^=0.68). There was no statistically significant effect of MC4R haploinsufficiency.

**Figure 8:**
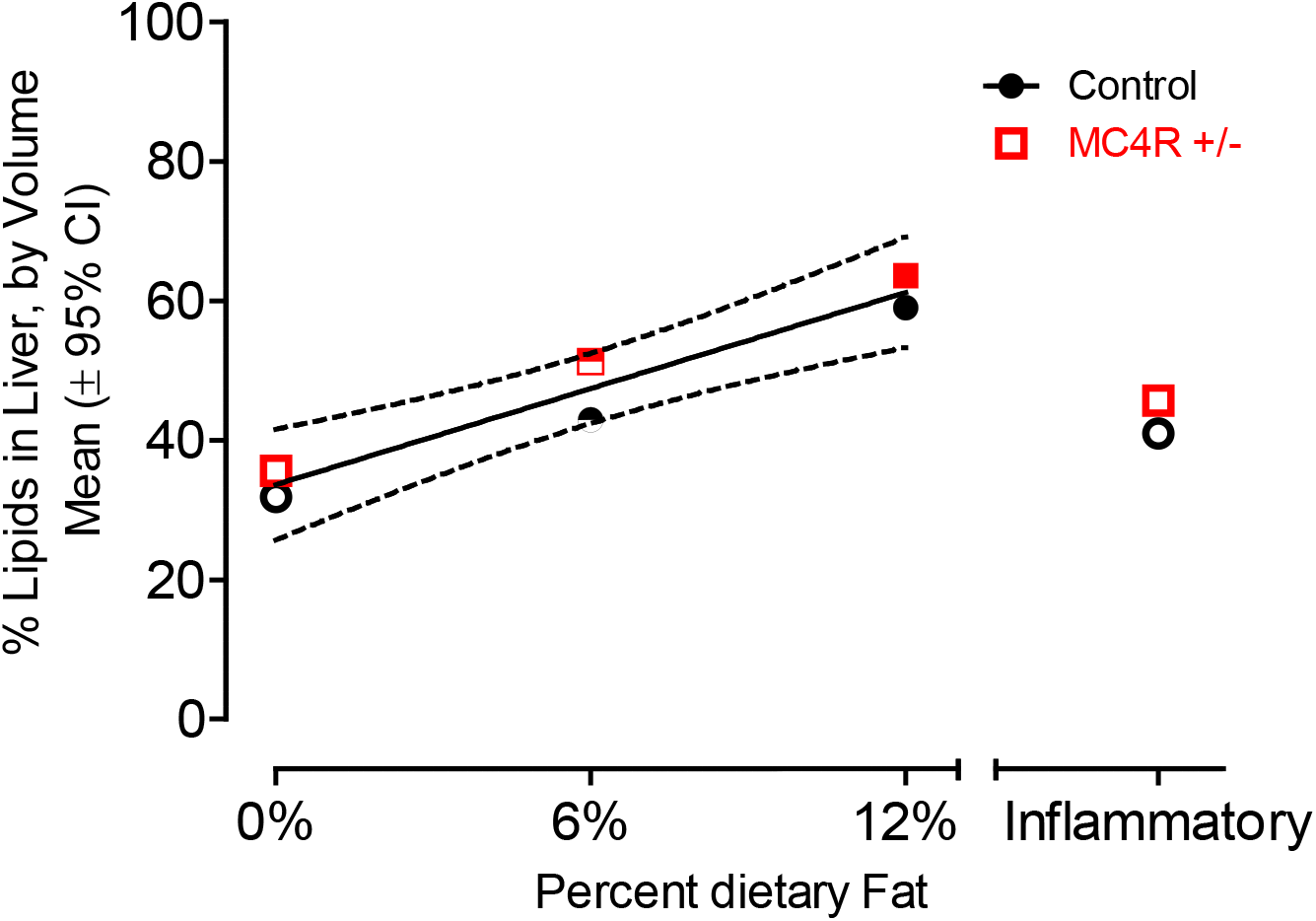
A clear linear relationship is observed between the percentage of dietary fat and the percentage of lipids in the liver (r^2^=0.68). The inflammation control group displayed values not significantly distinct from controls, indicating that this effect is not due to inflammation. No significant effect of MC4R haploinsufficiency was detected.

## 4. Discussion

The MC4R haploinsufficient rat displayed a phenotypic expression of obesity, consistent with the mutation of the MC4R receptor as the most common human monogenic cause of obesity. Motivational changes in the MC4R haploinsufficient rat, however, were intricately determined by two other major factors: age and diet. Prior to the onset of obesity, the MC4R +/- animals fed the control diet displayed an increased motivation to work for sucrose rewards on all measures of performance. However, consumption of the high saturated fat diet masked the effect of the MC4R haploinsufficiency. After obesity was well established, the MC4R +/- animals fed the control diet no longer showed the increased motivation to work for sucrose rewards, and in the presence of distracting stimuli, displayed a prominent decrease in motivation to work for sucrose rewards. In contrast, the MC4R +/- animals fed the high saturated fat diet showed increased motivation for reward, regardless of the value of the reward (shown by the variable progressive-ratio task), to help maintain their already rewarding dietary consumption habits. Furthermore, under the high saturated fat diet the MC4R +/- rats displayed significant resistance to the otherwise disrupting effect of the auditory tone distracting stimulus, earning significantly more rewards than the Wistar controls.

An enhancement of progressive-ratio performance for standard chow or sucrose is found in the Zucker obese rat relative to lean controls, in short-term (1-4 weeks) diet-induced obese rats and with the discontinuation of a high fat/high sugar diet in obesity prone rats (la Fleur *et al*., 2007; Rasmussen & Huskinson, 2008; Pickering *et al*., 2009); quite reminiscent of Epstein’s increased reinforcement with overweight vs. lean humans (Saelens & Epstein, 1996; Epstein *et al*., 2007).

Reward-related dopaminergic system areas of the brain are highly connected with the melanocortin system. POMC and AGRP neurons from the arcuate nucleus of the hypothalamus have projections to areas such as the ventral tegmental area, the nucleus accumbens, as well as the lateral hypothalamus (King & Hentges, 2011: Bagnol *et al*., 1999: Cui *et al*., 2012). Although the melanocortinergic and dopaminergic systems normally interact with their own receptors, there is growing evidence indicating that they may cross interact with the receptors of each other, at least in some brain regions implicated in feeding behaviors and motivation (He *et al*., 2015). For example, D1 and D2 receptors are co-localized with MC4R in the striatum and nucleus accumbens. Cui and colleagues identified a role for MC4R signaling in D1R neurons in learning both food-reinforced and non-food-reinforced procedural memories; in contrast, they failed to identify a role for MC4R signaling in the motivation to obtain palatable food (Cui *et al*., 2012). Both MC4R and D2 receptors work cohesively inside the bed nucleus of the stria terminalis (Yoon & Baik, 2015). Within the ventral tegmental area, it has been shown that injections of melanocortin receptor agonists have decreased consumption of palatable rewarding sucrose solutions during a two-bottle sucrose preference task (Yen & Roseberry, 2013). These studies demonstrate that melanocortins can act directly in the VTA to control reward-related feeding. Thus, these studies add to the growing body of evidence showing that melanocortins can interact with the mesolimbic dopamine system to regulate multiple reward-related behaviors. Along with the connection to dopamine, previous studies have linked motivational differences between control and MC4R haploinsufficient groups (Vaughan *et al*., 2006; Cui *et al*., 2013). The studies used both a progressive ratio and a fixed ratio respectively with motivational differences uncovered; however, age and dietary differences were not observed. A key difference between the previously cited studies and ours was the accessibility to food. Both studies use a form of food restriction while our animals had *ad libitum* access to their food. Availability of food, specifically with differing levels of saturated fat, further emphasizes the complex connection between the MC4R and dopamine reward systems. The complex relationship generalizes more uniquely to human individuals with MC4R deficits that have an abundant availability of easily accessible food.

A linear relationship was found between the percentage of dietary fat with the percentage of lipid deposits in the liver. The results indicate that both groups of animals have around 30% volume of fat in their livers on the control diet, and that ratio increases by 10-15% per 6% saturated fat added to the dietary condition. Control animals seem to have an above-average volume of lipid deposits; however, even the animals on the control diet had access to food *ab libitum*. The constant access to food might have increased their base level of fat in the liver especially compared to humans when food is not necessarily available at all times (i.e. while working, school, or simply following a normal three-meal diet). These results coincide with previous findings on the effect of dietary fat creating a similar representation of steatosis in our animals as they did with theirs (Ahmed *et al*., 2009).

Diet and MC4R+/- haploinsufficiency promoted prominent morphological changes in dendritic spine morphology, whereby a population shift towards increased dendritic spine length and decreased dendritic spine head diameter were observed in MC4R+/- haploinsufficient rats fed the high saturated diet relative to control animals. Fundamentally, the morphological shift towards a more immature dendritic spine phenotype in the MC4R+/- haploinsufficient rats fed the high-fat diet (i.e., thin vs. mushroom), supports alterations in synaptic function and efficacy. Indeed, dendritic spine length is negatively associated with synaptic efficacy (Araya *et al*., 2014) and dendritic spine head diameter is positively correlated with synaptic area (i.e., postsynaptic density; e.g., Harris & Stevens, 1988; 1989; Arellano *et al*., 2007). Synaptic area, in turn, is significantly associated with both the number of presynaptic (Harris & Stevens, 1988) and docked (Schikorski & Stevens, 1999) vesicles, as well as the number of postsynaptic receptors (Yuste, 2010). Taken together, the population shift towards increased dendritic spine length and decreased dendritic spine head diameter in MC4R+/- haploinsufficient rats fed the high saturated diet supports functional alterations in MSNs of the NAc; alterations which may underlie, at least in part, the observed motivational alterations.

Alterations to dendritic spine morphology are considered one of the hallmarks of neuroplasticity (Sala & Segal, 2014) and have been studied in chronic drug abuse models (Quintero, 2013; Gipson *et al*., 2013; Pal & Das, 2013). Cocaine-withdrawn animals present a marked increase in spine head diameter 45 minutes after cessation (Dumitriu *et al*., 2012; Toda *et al*., 2010) which may indicate a reversion in long-term potentiation capability. As substance abuse disorders are thought to physically manifest in reduced neuronal connectivity between the frontal cortex and basal ganglia (Motzkin *et al*., 2014), neuronal alterations in the NAc seen in this experiment may have both a causative and correlative effect on alterations to reward processing in the rat. Indeed, a high-fat diet in the rat has been shown to attenuate both motivation for sucrose reward as well as amphetamine-induced conditioned place preference (Davis *et al*., 2008), suggesting a mediating effect of dietary fat on dopaminergic turnover in the mesolimbic system. The focus of the present study was to pursue a better understanding of the interactions between overlapping reward and homeostatic neurocircuits for motivational systems with a focus on select neuroadaptations influencing dopaminergic neurotransmission in the central nervous system.

The results from the behavioral tasks, as well as dendritic spine morphology, indicate an exceptional role of motivation in the MC4R+/- haploinsufficient rat. During the early stages of the development of obesity, MC4R +/- animals that are not receiving an already rewarding high-fat diet display an increase in motivation towards food-related rewards. In the adult animals that have fully developed obesity, it seems that the maintenance of their obesity becomes the source of the motivational differences, causing the animals already fed the high saturated fat diet to display increased responding to food-related rewards. The signs of motivational differences during the early stages of obesity could indicate a dysregulation in reward pathways in the brain even prior to the development of obesity. While animals on different diets displayed motivational deficits at different time points during the development of obesity, an underlying dysregulation of the reward pathway could be the source. The knowledge of motivational differences caused by MC4R deficits reveals a potential new clinical target for the treatment of obesity in the underlying mechanisms of the dopamine reward circuitry connected to MC4R receptors.

## Abbreviations

MC4R: Melanocortin 4 receptor
POMC: Proopiomelanocortin
MSH: Melanocyte-stimulating hormone
AgRP: Agouti-related protein
FR: Fixed Ratio
PR: Progressive Ratio
MSNs: Medium spiny neurons

## 5. Acknowledgements

This work was supported in part by grants from NIH (National Institute on Drug Abuse, DA013137; National Institute of Child Health and Human Development, HD043680; National Institute of Mental Health, MH106392; National Institute of Neurological Disorders and Stroke, NS100624) and the interdisciplinary research training program supported by the University of South Carolina Behavioral-Biomedical Interface Program.

## References

Ahmed U, Redgrave TG, Oates PS. Effect of dietary fat to produce non-alcoholic fatty liver in the rat. J Gastroenterol Hepatol. 2009 Aug;24(8):1463–71. doi: 10.1111/j.1440-1746.2009.05870.x. PMID: 19702912.

Araya, R., Vogels, T. P., & Yuste, R. (2014). Activity-dependent dendritic spine neck changes are correlated with synaptic strength. Proc Natl Acad Sci U S A. 2014 Jul 15;111(28):E2895–904. doi: 10.1073/pnas.1321869111. PMID: 24982196; PMCID: PMC4104910.

Arellano, J. I., Benavides-Piccione, R., Defelipe, J., & Yuste, R. (2007). Ultrastructure of dendritic spines: correlation between synaptic and spine morphologies. Front Neurosci. 2007 Oct 15;1(1):131–43. doi: 10.3389/neuro.01.1.1.010.2007. PMID: 18982124; PMCID: PMC2518053.

Bagnol D, Lu XY, Kaelin CB, Day HE, Ollmann M, Gantz I, Akil H, Barsh GS, Watson SJ. Anatomy of an endogenous antagonist: Relationship between Agouti-related protein and proopiomelanocortin in brain. J Neurosci. 1999 Sep 15;19(18):RC26. doi: 10.1523/JNEUROSCI.19-18-j0004.1999. PMID: 10479719; PMCID: PMC6782481.

Barry D, Clarke M, Petry NM. Obesity and its relationship to addictions: Is overeating a form of addictive behavior? Am J Addict. 2009 Nov-Dec;18(6):439–51. doi: 10.3109/10550490903205579. PMID: 19874165; PMCID: PMC2910406.

Bae J, Sung BH, Cho IH, Kim SM, Song WK. NESH regulates dendritic spine morphology and synapse formation. PLoS One. 2012;7(4):e34677. doi: 10.1371/journal.pone.0034677. PMID: 22485184; PMCID: PMC3317636.

Blanpied TA, Ehlers MD. Microanatomy of dendritic spines: emerging principles of synaptic pathology in psychiatric and neurological disease. Biol Psychiatry. 2004 Jun 15;55(12):1121–7. doi: 10.1016/j.biopsych.2003.10.006. PMID: 15184030.

Casper RC, Sullivan EL, Tecott L. Relevance of animal models to human eating disorders and obesity. Psychopharmacology (Berl). 2008 Aug;199(3):313–29. doi: 10.1007/s00213-008-1102-2. PMID: 18317734.

Cone RD. Studies on the physiological functions of the melanocortin system. Endocr Rev. 2006 Dec;27(7):736–49. doi: 10.1210/er.2006-0034. PMID: 17077189.

Cui, H., Mason, B.L., Lee, C., Nishi, A., Elmquist, J.K., Lutter, M. Melanocortin 4 receptor signaling n dopamine 1 receptor neurons is required for procedural memory learning. Physiol Behav. 2012 May 15;106(2):201–10. doi: 10.1016/j.physbeh.2012.01.025. PMID: 22342812; PMCID: PMC3314089.

Cui H, Lutter M. The expression of MC4Rs in D1R neurons regulates food intake and locomotor sensitization to cocaine. Genes Brain Behav. 2013 Aug;12(6):658–65. doi: 10.1111/gbb.12057. PMID: 23786641; PMCID: PMC3782389.

Davis C, Strachan S, Berkson M. Sensitivity to reward: Implications for overeating and overweight. Appetite. 2004 Apr;42(2):131–8. doi: 10.1016/j.appet.2003.07.004. PMID: 15010176.

Davis JF, Tracy AL, Schurdak JD, Tschop MH, Lipton JW, Clegg DJ, Benoit SC. Exposure to elevated levels of dietary fat attenuates psychostimulant reward and mesolimbic dopamine turnover in the rat. Behav Neurosci. 2008 Dec;122(6):1257–63. doi: 10.1037/a0013111. PMID: 19045945; PMCID: PMC2597276.

de Leon J, Diaz FJ, Josiassen RC, Cooper TB, Simpson GM. Weight gain during a double-blind multidosage clozapine study. J Clin Psychopharmacol. 2007 Feb;27(1):22–7. doi: 10.1097/JCP.0b013e31802e513a. PMID: 17224708.

Dumitriu D, Laplant Q, Grossman YS, Dias C, Janssen WG, Russo SJ, Morrison JH, Nestler EJ. Subregional, dendritic compartment, and spine subtype specificity in cocaine regulation of dendritic spines in the nucleus accumbens. J Neurosci. 2012 May 16;32(20):6957–66. doi: 10.1523/JNEUROSCI.5718-11.2012. PMID: 22593064; PMCID: PMC3360066.

Epstein LH, Temple JL, Neaderhiser BJ, Salis RJ, Erbe RW, Leddy JJ. Food reinforcement, the dopamine D2 receptor genotype, and energy intake in obese and nonobese humans. Behav Neurosci. 2007 Oct;121(5):877–86. doi: 10.1037/0735-7044.121.5.877. Erratum in: Behav Neurosci. 2008 Feb;122(1):250. PMID: 17907820; PMCID: PMC2213752.

Farooqi IS, Keogh JM, Yeo GSH, Lank EJ, Cheetham T, O’Rahilly S. Clinical spectrum of obesity and mutations in the melanocortin 4 receptor gene. N Engl J Med. 2003 Mar 20;348(12):1085–95. doi: 10.1056/NEJMoa022050. PMID: 12646665.

Franco KN. Psychology and Psychiatry: Eating disorders. In: 2009 Current Clinical Medicine, Cleveland Clinic Foundation, 2009; ISBN: 978-1-4160-4096-5

Gipson CD, Reissner KJ, Kupchik YM, Smith AC, Stankeviciute N, Hensley-Simon ME, Kalivas PW. Reinstatement of nicotine seeking is mediated by glutamatergic plasticity. Proc Natl Acad Sci U S A. 2013 May 28;110(22):9124–9. doi: 10.1073/pnas.1220591110. PMID: 23671067; PMCID: PMC3670307.

Harris KM, Stevens JK. Dendritic spines of rat cerebellar Purkinje cells: serial electron microscopy with reference to their biophysical characteristics. J Neurosci. 1988 Dec;8(12):4455–69. doi: 10.1523/JNEUROSCI.08-12-04455.1988. PMID: 3199186; PMCID: PMC6569567.

Harris KM, Stevens JK. Dendritic spines of CA 1 pyramidal cells in the rat hippocampus: serial electron microscopy with reference to their biophysical characteristics. J Neurosci. 1989 Aug;9(8):2982–97. doi: 10.1523/JNEUROSCI.09-08-02982.1989. PMID: 2769375; PMCID: PMC6569708.

He ZG, Liu, BW, Xiang HB. Cross interaction of melanocortinergic and dopaminergic systems in neural modulation. Int J Physiol Pathophysiol Pharmacol. 2015 Dec 13;7(3):152-7. PMID: 26823964; PMCID: PMC4697671.

Ho G, MacKenzie RG. Functional characterization of mutations in melanocortin-4 receptor associated with human obesity. J Biol Chem. 1999 Dec 10;274(50):35816–22. doi: 10.1074/jbc.274.50.35816. PMID: 10585465.

Huszar D, Lynch CA, Fairchild-Huntress V, Dunmore JH, Fang Q, Berkemeier LR, Gu W, Kesterson RA, Boston BA, Cone RD, Smith FJ, Campfield LA, Burn P, Lee F. Targeted disruption of the melanocortin-4 receptor results in obesity in mice. Cell. 1997 Jan 10;88(1):131–41. doi: 10.1016/s0092-8674(00)81865-6. PMID: 9019399.

Jönsson EG, Nöthen MM, Grünhage F, Farde L, Nakashima Y, Propping P, Sedvall GC. Polymorphisms in the dopamine D2 receptor gene and their relationships to striatal dopamine receptor density of healthy volunteers. Mol Psychiatry. 1999 May;4(3):290–6. doi: 10.1038/sj.mp.4000532. PMID: 10395223.

Kenny PJ. Reward mechanisms in obesity: new insights and future directions. Neuron. 2011 Feb 24;69(4):664–79. doi: 10.1016/j.neuron.2011.02.016. PMID: 21338878; PMCID: PMC3057652.

Killeen PR, Posadas-Sanchez D, Johansen EB, Thrailkill EA. Progressive ratio schedules of reinforcement. J Exp Psychol Anim Behav Process. 2009 Jan;35(1):35–50. doi: 10.1037/a0012497. Erratum in: J Exp Psychol Anim Behav Process. 2009 Apr;35(2):152. PMID: 19159161; PMCID: PMC2806234.For

Kim DD, Basu A. Estimating the medical care costs of obesity in the United States: Systematic review, meta-analysis, and empirical analysis. Value Health. 2016 Jul-Aug;19(5):602–13. doi: 10.1016/j.jval.2016.02.008. PMID: 27565277.

King CM, Hentges ST. Relative number and distribution of murine hypothalamic proopiomelanocortin neurons innervating distinct target sites. PLoS One. 2011;6(10):e25864. doi: 10.1371/journal.pone.0025864. PMID: 21991375; PMCID: PMC3186811.

Krashes MJ, Lowell BB, Garfield AS. Melanocortin-4 receptor-regulated energy homeostasis. Nat Neurosci. 2016 Feb;19(2):206–19. doi: 10.1038/nn.4202. PMID: 26814590; PMCID: PMC5244821.

la Fleur SE, Vanderschuren LJ, Luijendijk MC, Kloeze BM, Tiesjema B, Adan RA. A reciprocal interaction between food-motivated behavior and diet-induced obesity. Int J Obes (Lond). 2007 Aug;31(8):1286–94. doi: 10.1038/sj.ijo.0803570. PMID: 17325683.

Leddy JJ, Epstein LH, Jaroni JL, Roemmich JN, Paluch RA, Goldfield GS, Lerman C. Influence of methylphenidate on eating in obese men. Obes Res. 2004 Feb;12(2):224–32. doi: 10.1038/oby.2004.29. PMID: 14981214.

Lieber CS, Leo MA, Mak KM, Xu Y, Cao Q, Ren C, Ponommarenko A, DeCarli LM. Model of nonalcoholic steatohepatitis. Am J Clin Nutr. 2004 Mar;79(3):502–9. doi: 10.1093/ajcn/79.3.502. PMID: 14985228.

Loos RJ, Lindgren CM, Li S, Wheeler E, Zhao JH, Prokopenko I, Inouye M, Freathy RM, Attwood AP, Beckmann JS, et al. Common variants near MC4R are associated with fat mass, weight and risk of obesity. Nat Genet. 2008 Jun;40(6):768–75. doi: 10.1038/ng.140. PMID: 18454148; PMCID: PMC2669167.

Lubrano-Berthelier C, Cavazos M, Le Stunff C, Haas K, Shapiro A, Zhang S, Bougneres P, Vaisse C. The human MC4R promoter: Characterization and role in obesity. Diabetes. 2003 Dec;52(12):2996–3000. doi: 10.2337/diabetes.52.12.2996. PMID: 14633862.

McLaurin KA, Fairchild AJ, Shi D, Booze RM, Mactutus CF. Valid statistical approaches for clustered data: A Monte Carlo simulation study. bioRxiv 2020.11.27.400945; doi: https://doi.org/10.1101/2020.11.27.400945.

McLaurin KA, Mactutus CF. Polytocus focus: Uterine position effect is dependent upon horn size. Int J Dev Neurosci. 2015 Feb;40:85–91. doi: 10.1016/j.ijdevneu.2014.11.001. PMID: 25447787; PMCID: PMC4451055.

Merino-Serrais P, Benavides-Piccione R, Blazquez-Llorca L, Kastanauskaite A, Rabano A, Avila J, DeFelipe J. The influence of phospho-tau on dendritic spines of cortical pyramidal neurons in patients with Alzheimer’s disease. Brain. 2013 Jun;136(Pt 6):1913–28. doi: 10.1093/brain/awt088. PMID: 23715095; PMCID: PMC3673457.

Motzkin JC, Baskin-Sommers A, Newman JP, Kiehl KA, Koenigs M. Neural correlates of substance abuse: Reduced functional connectivity between areas underlying reward and cognitive control. Hum Brain Mapp. 2014 Sep;35(9):4282–92. doi: 10.1002/hbm.22474. PMID: 24510765; PMCID: PMC4107096.

Mul JD, van Boxtel R, Bergen DJ, Brans MA, Brakkee JH, Toonen PW, Garner KM, Adan RA, Cuppen E. Melanocortin receptor 4 deficiency affects body weight regulation, grooming behavior, and substrate preference in the rat. Obesity (Silver Spring). 2012 Mar;20(3):612–21. doi: 10.1038/oby.2011.81. Erratum in: Obesity (Silver Spring). 2012 Jul;20(7):1545. Dosage error in article text. PMID: 21527895; PMCID: PMC3286758.

Novelli EL, Diniz YS, Galhardi CM, Ebaid GM, Rodrigues HG, Mani F, Fernandes AA, Cicogna AC, Novelli Filho JL. Anthropometrical parameters and markers of obesity in rats. Lab Anim. 2007 Jan;41(1):111–9. doi: 10.1258/002367707779399518. PMID: 17234057.

Ogden CL, Carroll MD, Lawman HG, Fryar CD, Kruszon-Moran D, Kit BK, Flegal KM. Trends in obesity prevalence among children and adolescents in the United States, 1988-1994 through 2013-2014. JAMA. 2016 Jun 7;315(21):2292–9. doi: 10.1001/jama.2016.6361. PMID: 27272581; PMCID: PMC6361521.

Pal A, Das S. Chronic morphine exposure and its abstinence alters dendritic spine morphology and upregulates Shank1. Neurochem Int. 2013 Jun;62(7):956–64. doi: 10.1016/j.neuint.2013.03.011. PMID: 23538264.

Paxinos, G., and Watson, C. (2014). The rat brain in stereotaxic coordinates, 7th ed. (Elsevier Academic Press).

Pickering C, Alsiö J, Hulting AL, Schiöth HB. Withdrawal from free-choice high-fat high-sugar diet induces craving only in obesity-prone animals. Psychopharmacology (Berl). 2009 Jun;204(3):431–43. doi: 10.1007/s00213-009-1474-y. PMID: 19205668.

Pijl H. Reduced dopaminergic tone in hypothalamic neural circuits: expression of a “thrifty” genotype underlying the metabolic syndrome? Eur J Pharmacol. 2003 Nov 7;480(1-3):125–31. doi: 10.1016/j.ejphar.2003.08.100. PMID: 14623356.

Pohjalainen T, Rinne JO, Någren K, Lehikoinen P, Anttila K, Syvälahti EK, Hietala J. The A1 allele of the human D2 dopamine receptor gene predicts low D2 receptor availability in healthy volunteers. Mol Psychiatry. 1998 May;3(3):256–60. doi: 10.1038/sj.mp.4000350. PMID: 9672901.

Preston RJ, Bishop GA, Kitai ST. Medium spiny neuron projection from the rat striatum: an intracellular horseradish peroxidase study. Brain Res. 1980 Feb 10;183(2):253–63. doi: 10.1016/0006-8993(80)90462-x. PMID: 7353139.

Quintero GC. Role of nucleus accumbens glutamatergic plasticity in drug addiction. Neuropsychiatr Dis Treat. 2013;9:1499–512. doi: 10.2147/NDT.S45963. PMID: 24109187; PMCID: PMC3792955.

Rasmussen EB, Huskinson SL. Effects of rimonabant on behavior maintained by progressive ratio schedules of sucrose reinforcement in obese Zucker (fa/fa) rats. Behav Pharmacol. 2008 Oct;19(7):735–42. doi: 10.1097/FBP.0b013e3283123cc2. PMID: 18797250.

Richardson NR, Roberts DC. Progressive ratio schedules in drug self-administration studies in rats: a method to evaluate reinforcing efficacy. J Neurosci Methods. 1996 May;66(1):1–11. doi: 10.1016/0165-0270(95)00153-0. PMID: 8794935.

Saelens BE, Epstein LH. Reinforcing value of food in obese and non-obese women. Appetite. 1996 Aug;27(1):41–50. doi: 10.1006/appe.1996.0032. PMID: 8879418.

Sala C, Segal M. Dendritic spines: the locus of structural and functional plasticity. Physiol Rev. 2014 Jan;94(1):141–88. doi: 10.1152/physrev.00012.2013. PMID: 24382885.

Schikorski T, Stevens CF. Quantitative fine-structural analysis of olfactory cortical synapses. Proc Natl Acad Sci U S A. 1999 Mar 30;96(7):4107–12. doi: 10.1073/pnas.96.7.4107. PMID: 10097171; PMCID: PMC22428.

Small DM, Jones-Gotman M, Dagher A. Feeding-induced dopamine release in dorsal striatum correlates with meal pleasantness ratings in healthy human volunteers. Neuroimage. 2003 Aug;19(4):1709–15. doi: 10.1016/s1053-8119(03)00253-2. PMID: 12948725.

Speakman J, Hambly C, Mitchell S, Król E. The contribution of animal models of obesity. Lab Anim. 2008 Oct;42(4):413–32. doi: 10.1258/la.2007.006067. PMID: 18782824.

Stice E, Spoor S, Bohon C, Small DM. Relation between obesity and blunted striatal response to food is moderated by TaqIA A1 allele. Science. 2008 Oct 17;322(5900):449–52. doi: 10.1126/science.1161550. PMID: 18927395; PMCID: PMC2681095.

Thomas JT, Brownell KD. Obesity. In: Ayers A, Baum A, McManus C, Newman S, Walston K, Weinman J, and West R. 2nd ed. Cambridge Handbook of Psychology, Health, and Medicine. Cambridge University Press: New York. 2007, pp. 797–800.

Toda S, Shen H, Kalivas PW. Inhibition of actin polymerization prevents cocaine-induced changes in spine morphology in the nucleus accumbens. Neurotox Res. 2010 Nov;18(3-4):410–5. doi: 10.1007/s12640-010-9193-z. PMID: 20401643.

Vaisse C, Clement K, Durand E, Hercberg S, Guy-Grand B, Froguel P. Melanocortin-4 receptor mutations are a frequent and heterogeneous cause of morbid obesity. J Clin Invest. 2000 Jul;106(2):253–62. doi: 10.1172/JCI9238. PMID: 10903341; PMCID: PMC314306.

Vaughan C, Moore M, Haskell-Luevano C, Rowland NE. Food-motivated behavior of melanocortin-4 receptor knockout mice under a progressive ratio schedule. Peptides. 2006 Nov;27(11):2829–35. doi: 10.1016/j.peptides.2006.07.008. PMID: 16930774.

Volkow ND, Wang GJ, Baler RD. Reward, dopamine and the control of food intake: implications for obesity. Trends Cogn Sci. 2011 Jan;15(1):37–46. doi: 10.1016/j.tics.2010.11.001. PMID: 21109477; PMCID: PMC3124340.

Volkow ND, Wang GJ, Fowler JS, Telang F. Overlapping neuronal circuits in addiction and obesity: Evidence of systems pathology. Philos Trans R Soc Lond B Biol Sci. 2008a Oct 12;363(1507):3191–200. doi: 10.1098/rstb.2008.0107. PMID: 18640912; PMCID: PMC2607335.

Volkow ND, Wang GJ, Telang F, Fowler JS, Thanos PK, Logan J, Alexoff D, Ding YS, Wong C, Ma Y, Pradhan K. Low dopamine striatal D2 receptors are associated with prefrontal metabolism in obese subjects: possible contributing factors. Neuroimage. 2008b Oct 1;42(4):1537–43. doi: 10.1016/j.neuroimage.2008.06.002. PMID: 18598772; PMCID: PMC2659013.

Wang GJ, Volkow ND, Logan J, Pappas NR, Wong CT, Zhu W, Netusil N, Fowler JS. Brain dopamine and obesity. Lancet. 2001 Feb 3;357(9253):354–7. doi: 10.1016/s0140-6736(00)03643-6. PMID: 11210998.

Wang GJ, Volkow ND, Thanos PK, Fowler JS. Positron emission tomography evidence of similarity between obesity and drug addiction. Psychiatric Annals. 2003; 33(2): 104–111. https://doi.org/10.3928/0048-5713-20030201-06

Wang GJ, Volkow ND, Thanos PK, Fowler JS. Similarity between obesity and drug addiction as assessed by neurofunctional imaging: a concept review. J Addict Dis. 2004;23(3):39–53. doi: 10.1300/J069v23n03_04. PMID: 15256343.

Wang Y, Beydoun MA, Min J, Xue H, Kaminsky LA, Cheskin LJ. Has the prevalence of overweight, obesity and central obesity leveled off in the United States? Trends, patterns, disparities, and future projections for the obesity epidemic. Int J Epidemiol. 2020 Jun 1;49(3):810–823. doi: 10.1093/ije/dyz273. PMID: 32016289; PMCID: PMC7394965. Fo

West MJ. Introduction to Stereology. Cold Spring Harb Protoc, 2012(8): pdb.top070623.

Wise RA. Dopamine and reward: The anhedonia hypothesis 30 years on. Neurotox Res. 2008 Oct;14(2-3):169–83. doi: 10.1007/BF03033808. PMID: 19073424; PMCID: PMC3155128.

Yen HH, Roseberry AG. Decreased consumption of rewarding sucrose solutions after injection of melanocortins into the ventral tegmental area of rats. Psychopharmacology(Berl). 2015; 232: 285–294.

Yoon YR, Baik JH. Melanocortin 4 receptor and dopamine D2 receptor expression in brain areas involved in food intake. Endocrinol Metab (Seoul). 2015; 30(4), 576–583.

Yuste R. (2010). Dendritic Spines. Cambridge, MA: Massachusetts Institute of Technology.

